# PHi-C2: interpreting Hi-C data as the dynamic 3D genome state

**DOI:** 10.1101/2022.05.06.490994

**Authors:** Soya Shinkai, Hiroya Itoga, Koji Kyoda, Shuichi Onami

## Abstract

Hi-C is a widely used assay for studying three-dimensional (3D) genome organization across the whole genome. Here, we present PHi-C2, a Python package supported by mathematical and biophysical polymer modeling, that converts an input Hi-C matrix data into the polymer model’s dynamics, structural conformations, and rheological features. The updated optimization algorithm to regenerate a highly similar Hi-C matrix provides a fast and accurate optimal solution compared to the previous version by eliminating a computational bottleneck in the iterative optimization process. Besides, we newly set up the availability on Google Colab workflow to run, easily change parameters and check the results in the notebook. Overall, PHi-C2 can be a valuable tool to mine the dynamic 3D genome state embedded in Hi-C data.

**Availability and Implementation:** PHi-C2 as the `phic` Python package is freely available under the GPL license and can be installed from the Python package index. The source code is available from GitHub at https://github.com/soyashinkai/PHi-C2. Without preparing a Python environment, PHi-C2 can run on Google Colab (https://bit.ly/3rlptGI).

## 1 Introduction

High-throughput chromosome conformation capture (Hi-C) quantifies genomic DNA contacts in the three-dimensional (3D) conformation of chromosomes across the whole genome (Lieberman-Aiden *et al*., 2009). The processed data are typically combined into a matrix as the population-averaged contact probability, which is depicted as a two-dimensional (2D) heatmap (Kerpedjiev *et al*., 2018; Robinson *et al*., 2018). The various 2D Hi-C patterns should reflect the structural characteristics of the 3D genome organization. However, since Hi-C data is nothing but a welter of snapshots of proximal genomic DNA pairs due to chemical fixation, the outcome pictures are mostly limited as static and averaged models. Meanwhile, live-cell imaging has revealed that chromatin dynamically moves coupling with genome functions within living cells (Heun *et al*., 2001; Nagashima *et al*., 2019). Therefore, biophysical modeling is essential toward a quantitative understanding of the gap between Hi-C data for fixed cells and chromatin dynamics information for living cells.

In 2020, we released the PHi-C software as Python codes designed to decipher Hi-C data into polymer dynamics (Shinkai *et al*., 2020b). PHi-C demonstrations output dynamic characteristics of genomic loci and chromosomes, as observed in live-cell imaging experiments, and allow for interpreting Hi-C data as the dynamic information of the 3D genome organization (Shinkai *et al*., 2020a). However, although the reconstructed contact matrix from an input contact matrix is in excellent agreement with the Pearson’s correlation of more than 95% (Shinkai *et al*., 2020c), the optimization procedure, which is a core part of the PHi-C algorithm, is a computational bottleneck; the iterative algorithm to reduce the cost function at every optimization step practically takes a few days to obtain an optimal solution. Furthermore, due to defining the cost function by the logarithmic form in the optimization and interpolating for the null value of an input contact matrix, PHi-C is not appropriate for every Hi-C matrix data.

To overcome these problems, first, we found the mathematical transformation between an input contact matrix and a set of parameters of our polymer model; the forward and inverse transformations are in the invertible correspondence (Supplementary Note S1). Then, we elucidated the mathematical concept of the optimization and updated the optimization algorithm (Supplementary Note S2). Benchmarks of the optimization calculations indicated improved scores for both speed and the closeness between the input and optimal contact matrices (Supplementary Note S3). In addition, we incorporated the rheology analysis (Shinkai *et al*., 2020a) as a new function to convert Hi-C data into the spectrum of the dynamic rheological properties along the genomic coordinate of a single chromosome (Supplementary Note S4). Here, we present PHi-C2 as a Python package, which has been redesigned from the ground up with additional new features. Additionally, we shipped a command-line interface (CLI) for convenient application.

## 2 Implementation and benchmarks

PHi-C2 is the `phic` Python package, which inclues a suite of CLI subcommands under a top-level `phic` command namespace (Fig. 1). The input Hi-C file is the contact matrix format extracted from the .hic file by Juicer and Straw (Durand *et al*., 2016). First, the `phic preprocessing` command converts the input into the normalized contact matrix data so that the diagonal elements are units based on the PHi-C polymer modeling theory. Next, the `phic optimization` command outputs an optimal matrix as the PHi-C polymer model parameter set. To visualize the results of a reconstructed contact matrix and a contact probability decay curve, we prepared the `phic plot-optimization` command. After the optimization procedure, users can calculate the polymer model’s dynamics and structure sampling using the `phic dynamics` and `phic sampling` commands, respectively. The outputs are .xyz and .psf format files; visualization requires VMD (Humphrey *et al*., 1996). Furthermore, to reveal the hierarchical and dynamic 3D genome state embedded in the input 2D Hi-C pattern, users can apply the `phic rheology` command and the three consecutive commands (`phic plot–compliance, phic plot-modulus`, and `phic plot-tangent`) to visualize the rheological analysis results. In addition, without introducing a Python environment in the user’s local platform, PHi-C2 runs on Google Colab; users can easily change parameters and check the results of the plots along the workflow in the notebook.

**Figure 1.**
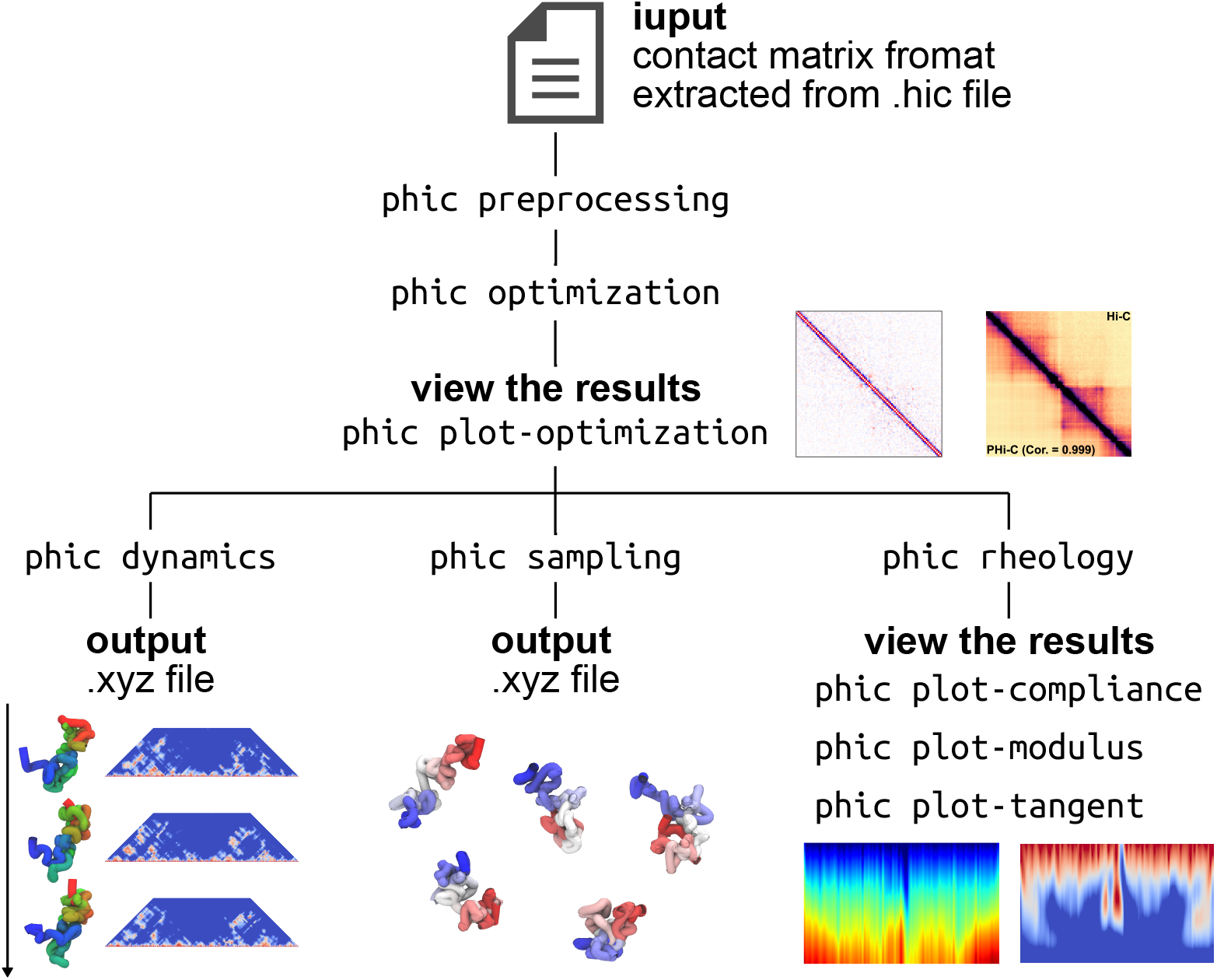
Overview of the PHi-C2 pipeline and `phic` CLI commands. PHi-C2 analyzes input Hi-C data as the contact matrix format extracted from.hic file by Juicer and Straw (Durand *et al*., 2016). The updated optimization algorithm outputs an optimal parameter set for the polymer model, and depicts the optimal contact matrix. Using the optimal parameters, users can calculate the polymer model’s dynamics, structure sampling, and rheology spectra consistent with the input Hi-C matrix.

As the optimization procedure is a core computational part of PHi-C2, we benchmarked the performance for a 400 × 400-sized input Hi-C matrix (Supplementary Note S3). First, the updated PHi-C2 algorithm improved the speed and accuracy compared to that obtained using the previous PHi-C version (Supplementary Figure S3). By varying the optimization parameters regarding the initial values, learning rate, and stop condition, we obtained optimal solutions with more than 99.7% agreement in 30 min (Supplementary Table S2). All tests were performed using an Intel®Xeon®Gold 6154 processor (24.75M Cache, 3.00 GHz) and Intel®distribution for the Python environment. The iterative optimization process depends on the input matrix size (Shinkai *et al*., 2020b), while 100 ×100–500 ×500 is a practically good input matrix size according to the genomic region of interest.

## 3 Conclusion

We developed a Python package, PHi-C2, to analyze Hi-C matrix data, including CLI subcommands for convenient manipulation. As we reconsidered the mathematical framework and eliminated the computational bottleneck of the previous version, the speed and accuracy improved. Therefore, without a massive computational cost, users can calculate the polymer dynamics, structural conformations, and rheological features consistent with the input Hi-C data. The easy installation from the Python package index and calculations on Google Colab would help users reveal the physical features embedded in Hi-C data.

## Acknowledgements

We thank Dr. M. Nakagawa for the critical feedback and helpful discussion provided.

