## Supplementary Information for "PHi-C2: interpreting Hi-C data as the dynamic 3D genome state"

### SUPPLEMENTARY INFORMATION
### PHi-C2: interpreting Hi-C data as the dynamic 3D genome state

**Soya Shinkai**[1], **Hiroya Itoga**[1], **Koji Kyoda**[1], **and Shuichi Onami**[1,2]

[1]Laboratory for Developmental Dynamics, RIKEN Center for Biosystems Dynamics Research, Kobe, 650-0047, Japan.
[2]Life Science Data Sharing Unit, Infrastructure Research and Development Division, RIKEN Information R&D and Strategy Headquarters, Kobe, 650-0047, Japan.

#### ABSTRACT

Supplementary information includes 4 Supplementary Notes with 5 Supplementary Figures and 2 Supplementary Tables.

#### Supplementary Note S1: Inverse transformation to reconstruct an input Hi-C matrix

PHi-C is a simulation tool to decipher Hi-C data into polymer dynamics based on the mathematical formalism of the polymer network model[1]. The normalized interaction matrix $\bar{\mathsf{K}} = \left(\bar{K}_{ij}\right)$ of the model has a one-to-one correspondence to a normalized contact matrix $\bar{\mathsf{C}} = \left(\bar{C}_{ij}\right)$ through a succession of matrix transformations. (i) $\bar{\mathsf{K}}$ into the normalized Laplacian matrix $\bar{\mathsf{L}} = \left(L_{ij}\right)$ by $\bar{\mathsf{L}} = \bar{\mathsf{D}} - \bar{\mathsf{K}}$, where the normalized degree matrix $\bar{\mathsf{D}} = \mathrm{diag}(\bar{D}_0, \bar{D}_1, ..., \bar{D}_{N-1})$, where $\bar{D}_i = \sum_{j=0}^{N-1} \bar{K}_{ij}$. (ii) As the Laplacian matrix $\bar{\mathsf{L}}$ is real symmetric, $\bar{\mathsf{L}}$ is diagonalizable. Besides, the $N$ eigenvalues satisfy $0 = \bar{\lambda}_0 < \bar{\lambda}_1 < \cdots < \bar{\lambda}_{N-1}$, as long as $\bar{\mathsf{L}}$ is positive semidefinite and there is an orthogonal matrix $\mathsf{Q}$ such that $\mathsf{Q}^{\mathsf{T}}\bar{\mathsf{L}}\mathsf{Q} = \mathrm{diag}(0, \bar{\lambda}_1, \cdots, \bar{\lambda}_{N-1})$. Then, $\bar{\mathsf{L}}$ is transformed into the normalized covariance matrix $\bar{\mathsf{M}} = \left(\bar{M}_{ij}\right)$ by $\bar{\mathsf{M}} = \mathsf{Q}\,\mathrm{diag}\left(0, \bar{\lambda}_1^{-1}, \cdots, \bar{\lambda}_{N-1}^{-1}\right)\mathsf{Q}^{\mathsf{T}}$. (iii) $\bar{\mathsf{M}}$ into the normalized variance matrix $\bar{\mathsf{\Sigma}}^2 = \left(\bar{\Sigma}_{ij}^2\right) = \left(\frac{\bar{M}_{ii} + \bar{M}_{jj} - 2\bar{M}_{ij}}{3}\right)$. (iv) $\bar{\mathsf{\Sigma}}^2$ into the contact matrix $\mathsf{C} = \left(1 + \bar{\mathsf{\Sigma}}^2\right)^{-3/2}$. Here, we briefly express this procedure as $\mathsf{C} = \varphi(\bar{\mathsf{K}})$. If we can find the inverse function $\varphi^{-1}$, we can solve the inverse problem, $\bar{\mathsf{K}} = \varphi^{-1}(\mathsf{C})$, for any input contact matrix.

The first step of the inverse function is $\bar{\mathsf{\Sigma}}^2 = \left(\mathsf{C}^{-2/3} - 1\right)$. As we noted[1], there is an inverse transformation from $\bar{\mathsf{\Sigma}}^2$ to $\bar{\mathsf{M}}$ because the matrix $\bar{\mathsf{M}}$ satisfies the condition $\bar{\mathsf{M}}\mathbf{1} = \mathbf{0}$, where $\mathbf{1}$ is a vector $(1, 1, \cdots, 1)^{\mathsf{T}}$. Here, we explicitly give the inverse transformation. Let $\boldsymbol{m}$ be a vector that consists of the diagonal elements of the matrix $\bar{\mathsf{M}}$, $\left(\bar{M}_{00}, \bar{M}_{11}, \cdots, \bar{M}_{N-1,N-1}\right)^{\mathsf{T}}$, and we define a square matrix $\mathsf{B}$ as $(\boldsymbol{m}, \boldsymbol{m}, \cdots, \boldsymbol{m})$. Then, the relationship between $\bar{\mathsf{\Sigma}}^2$ and $\bar{\mathsf{M}}$ can be written by $3\bar{\mathsf{\Sigma}}^2 = \mathsf{B} + \mathsf{B}^{\mathsf{T}} - 2\bar{\mathsf{M}}$. Multiplying $\mathbf{1}$ from right, we obtain $3\bar{\mathsf{\Sigma}}^2\mathbf{1} = \mathsf{A}\boldsymbol{m}$, where $\mathsf{A}$ is a square matrix defined by $N\mathsf{I} + \mathbf{1}\mathbf{1}^{\mathsf{T}}$. Since the matrix $\mathsf{A}$ is non-singular, there exists the inverse matrix given by $\mathsf{A}^{-1} = \frac{2N\mathsf{I} - \mathbf{1}\mathbf{1}^{\mathsf{T}}}{2N^2}$. Therefore, we can determine the diagonal elements of the matrix $\bar{\mathsf{M}}$ by $\boldsymbol{m} = 3\mathsf{A}^{-1}\bar{\mathsf{\Sigma}}^2\mathbf{1}$ and calculate all the elements by $\bar{M}_{ij} = \frac{\bar{M}_{ii} + \bar{M}_{jj} - 3\bar{\Sigma}_{ij}^2}{2}$.

Next, we considered a transformation from $\bar{\mathsf{M}}$ to $\bar{\mathsf{L}}$. Theoretically, the matrix $\bar{\mathsf{M}}$ should be positive semidefinite from the definition. However, $\bar{\mathsf{M}}$ calculated through the above transformation did not satisfy the positive semidefiniteness. Mainly, it would stem from the experimental noise of Hi-C data. Therefore, we could not find the Laplacian matrix $\bar{\mathsf{L}}$ straightforwardly by solving the eigenvalue problem. Instead, we took a different way to construct a pseudo-Laplacian matrix $\tilde{\mathsf{L}}$ using the Moore-Penrose (MP) inverse matrix $\bar{\mathsf{M}}^+$ of the matrix $\bar{\mathsf{M}}$[2]. As both matrices $\bar{\mathsf{M}}$ and $\bar{\mathsf{L}}$ theoretically satisfy the relationship $\bar{\mathsf{M}}\bar{\mathsf{L}} = \mathsf{I} - \frac{\mathbf{1}\mathbf{1}^{\mathsf{T}}}{N}$, the MP inverse matrix minimizes the Frobenius norm $\left\|\bar{\mathsf{M}}\bar{\mathsf{L}} - \mathsf{I} + \frac{\mathbf{1}\mathbf{1}^{\mathsf{T}}}{N}\right\|_{\mathrm{F}}$ and provides the solution by $\tilde{\mathsf{L}} = \bar{\mathsf{M}}^+\left(\mathsf{I} - \frac{\mathbf{1}\mathbf{1}^{\mathsf{T}}}{N}\right)$. The construction of the MP inverse matrix is based on the singular value decomposition of the matrix $\bar{\mathsf{M}}$, and we can easily implement it on numerical scripts. Finally, we can obtain a pseudo-interaction matrix $\tilde{\mathsf{K}}$ by $\tilde{\mathsf{K}} = -\tilde{\mathsf{L}}$ and zero-diagonals ($\tilde{K}_{ii} = 0$).

Based on the above algorithm, we reconstructed a contact matrix $\tilde{\mathsf{C}} = \varphi(\tilde{\mathsf{K}})$ from an input matrix $\mathsf{C}$ (Supplementary Figure S1A). Note that, although the pseudo-Laplacian matrix $\tilde{\mathsf{L}}$ is no longer positive semidefinite (Supplementary Figure S1B), the numerical calculation of the eigenvalues and eigenvectors works uneventfully. We estimated the closeness of all the elements between $\mathsf{C}$ and $\tilde{\mathsf{C}}$. Surprisingly, the `allclose` function in NumPy returned True, which means that two matrices are element-wise equal within the default numerical tolerance. Taken together, we found the inverse transformation $\varphi^{-1}$.

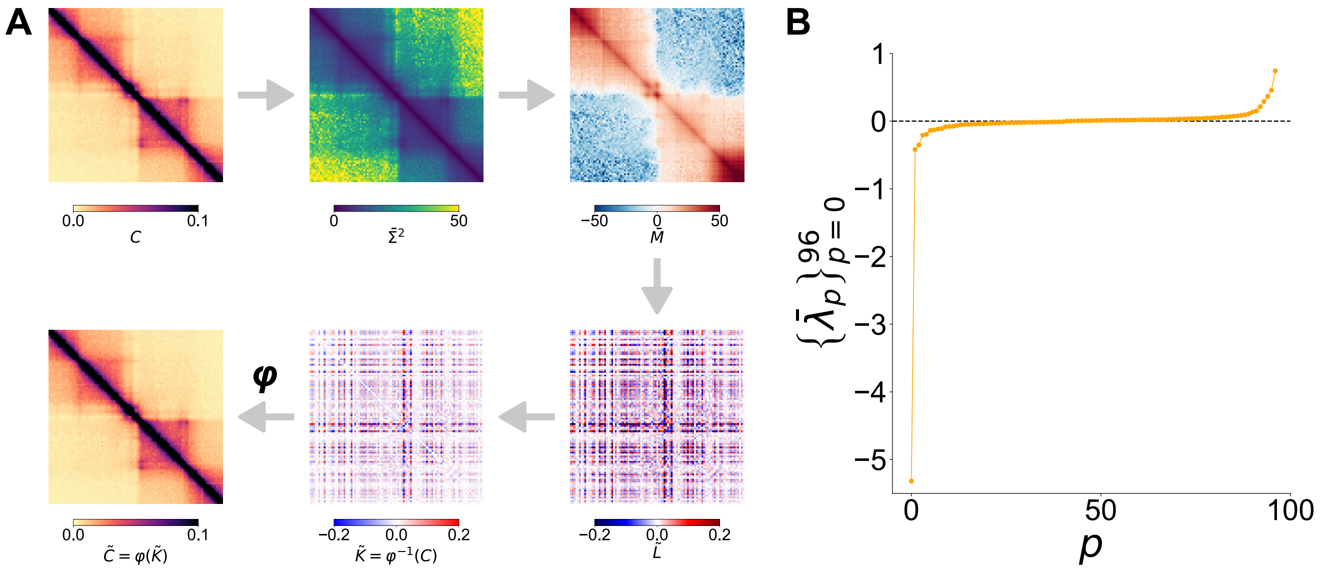


**Supplementary Figure S1. Matrix transformations in the PHi-C polymer modeling. (A)** The inverse transformation $\varphi^{-1}$ converts an input normalized contact matrix C [chr8: 42,100–44,525-kb (25-kb bins) for mouse embryonic stem cells (mESCs)[3]] into the pseudo-interaction matrix $\tilde{K}$ via matrices $\bar{\Sigma}^2$, $\bar{M}$ and $\tilde{L}$. The reconstructed contact matrix $\tilde{C} = \varphi(\tilde{K})$ and the input contact matrix C are equal within the default numerical tolerance. **(B)** Eigenvalues of the pseudo-Laplacian matrix $\tilde{L}$. Since there are negative eigenvalues, $\tilde{L}$ is no longer positive semidefinite.

#### Supplementary Note S2: Updated optimization operation

##### Mathematical concept of the PHi-C optimization

The optimization procedure is a central part of PHi-C. We define the optimization as an iterative finding of the normalized interaction matrix $\bar{\mathsf{K}}$ that decreases a cost function $f(\mathsf{C}, \varphi(\bar{\mathsf{K}}))$ for an input contact matrix $\mathsf{C}$. To clarify the mathematical concept, we introduce the following sets (Supplementary Figure S2):

- $\mathbb{K}^N = \left\{\bar{\mathsf{K}} | \text{ a symmetric } N \times N \text{ matrix with zero diagonals } \bar{K}_{ii} = 0\right\}$,
- $U_\epsilon = \left\{\bar{\mathsf{K}} \in \mathbb{K}^N | f(\mathsf{C}, \varphi(\bar{\mathsf{K}})) < \epsilon\right\}$,
- $A = \left\{\bar{\mathsf{K}} \in \mathbb{K}^N | \text{ the induced Laplacian matrix } \bar{\mathsf{L}} \text{ is positive-semidefinite and } \bar{K}_{i,i+1} > 0\right\}$.

For a small $\epsilon$, a matrix $\bar{\mathsf{K}} \in U_\epsilon$ is an optimal candidate. In Supplementary Note 1, we found the minimal solution $\varphi^{-1}(\mathsf{C}) = \tilde{\mathsf{K}} \in U_{\epsilon=0}$ by the inverse transformation. However, since the induced Laplacian matrix does not satisfy the positive semidefiniteness, the polymer system becomes unstable, and the normalized interaction matrix $\tilde{\mathsf{K}}$ is unrealistic. Thus, the set $A$ ensures the physically acceptable matrix $\bar{\mathsf{K}}$ with connections along the polymer backbone, $\bar{K}_{i,i+1} > 0$, as a stable polymer system.

Let $\bar{\mathsf{K}}_n$ be the normalized interaction matrix at the $n$-th iteration step in the optimization procedure. An initial matrix $\bar{\mathsf{K}}_0$ should be in the set $A$, and an optimization path in $\mathbb{K}^N$ would go into the set $U_\epsilon$ for a small $\epsilon$ (Supplementary Figure S2). Then, paths that output an optimal $\bar{\mathsf{K}} \in U_\epsilon \cap A$ are acceptable, but paths that go into $U_\epsilon \cap A^c$ should be forbidden.

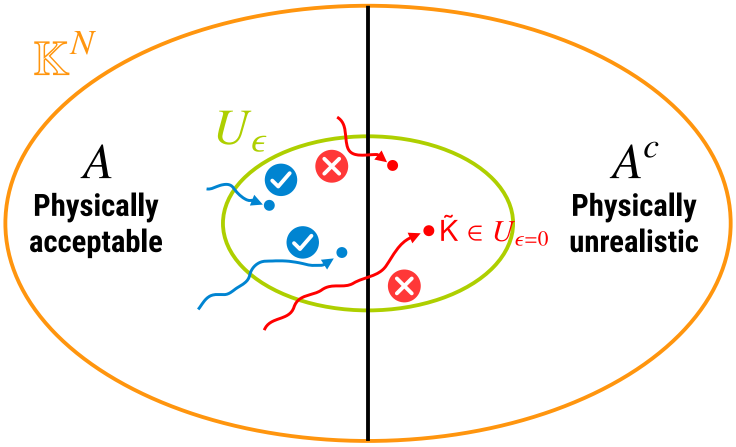


**Supplementary Figure S2. Mathematical concept of the PHi-C optimization**. The PHi-C optimization iteratively decreases the cost function in $\mathbb{K}^N$. The set $U_\epsilon$ has optimal candidates of the matrix $\bar{\mathsf{K}}$. As the inverse transformation induces the pseudo-interaction matrix $\tilde{\mathsf{K}}$, the matrix $\tilde{\mathsf{K}}$ is just an element of the set $U_{\epsilon=0}$. Optimization paths can go into the set $U_\epsilon \cap A^c$, where the polymer systems are physically unrealistic. Optimization paths should move in the physically acceptable set $A$.

##### Updated algorithm

We mainly updated the optimization algorithm regarding the definition of the cost function and the iterative operation at each step (Supplementary Table S1). Accordingly, both factors provided faster and more accurate optimization results than the previous version.

First, we re-defined the cost function as

$$f(\mathsf{C}, \varphi(\bar{\mathsf{K}})) = \sqrt{\frac{1}{N^2} \sum_{i=0}^{N-1} \sum_{j=0}^{N-1} \left[C_{ij} - (\varphi(\bar{\mathsf{K}}))_{ij}\right]^2}.$$

| | PHi-C1 | PHi-C2 |
|---|---|---|
| Cost function: $f(\mathsf{C},\varphi(\bar{\mathsf{K}}))$ | $\sqrt{\frac{1}{N^2}\sum_{i=0}^{N-1}\sum_{j=0}^{N-1}\left[\log_{10}C_{ij}-\log_{10}(\varphi(\bar{\mathsf{K}}))_{ij}\right]^2}$ | $\sqrt{\frac{1}{N^2}\sum_{i=0}^{N-1}\sum_{j=0}^{N-1}\left[C_{ij}-(\varphi(\bar{\mathsf{K}}))_{ij}\right]^2}$ |
| Optimization operation at every iteration step | For a randomly selected pair $(i,j)$,<br>$\xi = \eta \times \text{random.uniform}(0,1)$<br>**if** $\log_{10}(\varphi(\bar{\mathsf{K}}_{n-1}))_{ij} > \log_{10}C_{ij}$ **then**<br>$(\bar{\mathsf{K}}_n)_{ij} = (\bar{\mathsf{K}}_{n-1})_{ij} - \xi$<br>$(\bar{\mathsf{K}}_n)_{ji} = (\bar{\mathsf{K}}_{n-1})_{ji} - \xi$<br>**else**<br>$(\bar{\mathsf{K}}_n)_{ij} = (\bar{\mathsf{K}}_{n-1})_{ij} + \xi$<br>$(\bar{\mathsf{K}}_n)_{ji} = (\bar{\mathsf{K}}_{n-1})_{ji} + \xi$ | For all the elements,<br>$\bar{\mathsf{K}}_n = \bar{\mathsf{K}}_{n-1} - \eta\left(\varphi(\bar{\mathsf{K}}_{n-1}) - \mathsf{C}\right)$ |

**Supplementary Table S1. Mainly updated algorithms of PHi-C2 compared to PHi-C1.** $\eta$ is the learning rate parameter in the optimization operation.

This value represents the standard deviation between the contact probabilities $\mathsf{C}$ and $\varphi(\bar{\mathsf{K}})$ per element. On the other hand, we calculated the standard deviation between the logarithmic contact probabilities in the previous version (Supplementary Table S1). Since the logarithmic function is not defined for zero value, we needed an interpolate operation for the zero-valued elements so that the shape of the contact probability curve is unaltered[1]. Therefore, the newly defined cost function does not require the interpolation process.

Every iterative optimization step updates all the elements of the matrix $\bar{\mathsf{K}}_n$, obeying the following rule:

$$\bar{\mathsf{K}}_n = \bar{\mathsf{K}}_{n-1} - \eta\left(\varphi(\bar{\mathsf{K}}_{n-1}) - \mathsf{C}\right),$$

where $\eta$ is the learning rate parameter. In the previous PHi-C, the random choice of a matrix element $(i, j)$ and its slight change of the element value at every iteration step were a bottleneck (Supplementary Table S1). The new iterative rule overcomes the bottleneck by updating the values of all the elements at a time.

Lastly, we set a stop condition due to a monotonical decrease of the cost function,

$$0 < f(\mathsf{C},\varphi(\bar{\mathsf{K}}_{n-1})) - f(\mathsf{C},\varphi(\bar{\mathsf{K}}_n)) < \delta \quad \Longrightarrow \quad \text{output } \bar{\mathsf{K}}_n \text{ as an optimal solution } \bar{\mathsf{K}}_{\text{opt}},$$

where $\delta = \eta\alpha$ and $\alpha$ is a parameter for the stop condition.

Overall, the above three points are significant updates of the PHi-C2 algorithm. So far, we do not mathematically ensure that every optimization path decreases the cost function monotonically in the set $A$ (Supplementary Figure S2) since the update coefficient $\left(\varphi(\bar{\mathsf{K}}_{n-1}) - \mathsf{C}\right)$ does not exactly correspond to the gradient $\left.\frac{\partial f(\mathsf{C},\varphi(\bar{\mathsf{K}}))}{\partial \bar{K}_{ij}}\right|_{\bar{\mathsf{K}}=\bar{\mathsf{K}}_{n-1}}$, which has already been formulated[4]. However, we can operationally obtain an optimal solution in the set $A$ by tuning the initial point $\bar{\mathsf{K}}_0 \in A$, the learning rate $\eta$, and the stop condition parameter $\alpha$.

#### Supplementary Note S3: Benchmarks

After the optimization, we can estimate the closeness of the optimal contact matrix $\mathsf{C}_{\text{opt}} = \varphi(\bar{\mathsf{K}}_{\text{opt}})$ to the input contact matrix $\mathsf{C}$ using Pearson's correlation coefficient $r$ for the non-diagonal elements of both matrices. Here we show benchmarks for Hi-C data of mESCs[3] [chr1: 50–60 Mb (25-kb bins)].

For the $400 \times 400$-sized input matrix of mESCs and a default set of optimization parameters ($(\bar{\mathsf{K}}_0)_{i,i+1} = 0.5$, $\eta = 10^{-4}$, $\alpha = 10^{-4}$), PHi-C2 output an optimal solution with $r = 0.998$ in 26,307 iteration steps (Supplementary Figure S3A and Supplementary Table S2). On the other hand, for the same iteration steps and $\eta = 4 \times 10^{-3}$, an optimal solution of the previous PHi-C (PHi-C1) indicated $r = 0.966$ (Supplementary Figure S3B), although the meaning of the cost function and the learning rate differ between PHi-C1 and PHi-C2. We can confirm that the updated algorithm improves the speed and accuracy of the optimization procedure compared to PHi-C1.

Next, we show benchmarks for different optimization parameters: initial values of the polymer backbone $(\bar{\mathsf{K}}_0)_{i,i+1}$, the learning rate $\eta$, and the stop condition parameter $\alpha$ (Supplementary Tabel S2). We carried out all the calculations using Intel® Xeon® Gold 6154 processor (24.75M Cache, 3.00 GHz) and Intel® distribution for Python environment. The procedure stopped within less than 100,000 iteration steps and 30 min for these parameters. Interestingly, the optimal cost function values differently varied, but the correlation values almost showed 0.998. As our stop condition relates to the cost function curve saturation, the optimal solutions would be close to local minimums in the set $U_{\varepsilon=0.00223}$. Also, we could confirm that the optimal solutions are located in the set $A$ due to the positive semidefiniteness (Supplementary Figure S3C).

As we stated, the update algorithm does not always decrease the cost function monotonically. We observed such counter-examples for Hi-C data of the mouse erythroid cells[5] [chr1: 3–195.5 Mb (500-kb bins)] in the prometaphase. Here, we altered only the initial values of the polymer backbone $(\bar{\mathsf{K}}_0)_{i,i+1} = 0.1 \sim 1.0$ (fixed $\eta = 10^{-4}$ and $\alpha = 10^{-4}$). Some curves showed an increase in the cost function within the initial 100 steps (Supplementary Figure S4A). Nevertheless, finally, every curve converged and output an optimal solution. On the other hand, for Hi-C data in the late G1 phase, we confirmed a monotonical decrease of the cost function (Supplementary Figure S4B).

| $(\bar{\mathsf{K}}_0)_{i,i+1}$ | $\eta$ | $\alpha$ | # iterations | time | $f(\mathsf{C}, \varphi(\bar{\mathsf{K}}_{\text{opt}}))$ [$\times 10^{-3}$] | $r$ |
|---|---|---|---|---|---|---|
| 0.1 | $10^{-4}$ | $10^{-4}$ | 68,164 | 24 m 27 s | 2.10086 | 0.998 |
| 0.2 | $10^{-4}$ | $10^{-4}$ | 57,477 | 20 m 52 s | 2.09129 | 0.998 |
| 0.3 | $10^{-4}$ | $10^{-4}$ | 36,388 | 13 m 21 s | 2.11411 | 0.998 |
| 0.4 | $10^{-4}$ | $10^{-4}$ | 17,931 | 7 m 00 s | 1.94762 | 0.997 |
| 0.5 | $10^{-4}$ | $10^{-4}$ | 26,307 | 9 m 54 s | 1.73502 | 0.998 |
| 0.6 | $10^{-4}$ | $10^{-4}$ | 35,097 | 13 m 48 s | 1.62454 | 0.998 |
| 0.7 | $10^{-4}$ | $10^{-4}$ | 43,714 | 16 m 51 s | 1.56127 | 0.998 |
| 0.8 | $10^{-4}$ | $10^{-4}$ | 52,089 | 19 m 54 s | 1.52338 | 0.998 |
| 0.9 | $10^{-4}$ | $10^{-4}$ | 60,228 | 20 m 48 s | 1.50057 | 0.998 |
| 1.0 | $10^{-4}$ | $10^{-4}$ | 68,139 | 24 m 34 s | 1.48751 | 0.998 |
| 0.5 | $2 \times 10^{-4}$ | $10^{-4}$ | 13,154 | 4 m 24 s | 1.73501 | 0.998 |
| 0.5 | $10^{-4}$ | $10^{-4}$ | 26,307 | 9 m 54 s | 1.73502 | 0.998 |
| 0.5 | $5 \times 10^{-5}$ | $10^{-4}$ | 52,613 | 17 m 35 s | 1.73502 | 0.998 |
| 0.5 | $10^{-4}$ | $10^{-3}$ | 13,369 | 4 m 31 s | 2.22941 | 0.997 |
| 0.5 | $10^{-4}$ | $10^{-4}$ | 26,307 | 9 m 54 s | 1.73502 | 0.998 |
| 0.5 | $10^{-4}$ | $10^{-5}$ | 42,895 | 14 m 15 s | 1.67631 | 0.998 |

**Supplementary Table S2. Benchmarks of PHi-C2 for different optimization parameters.** We used Hi-C data of mESCs [chr1: 50–60 Mb (25-kb bins)] as an input.

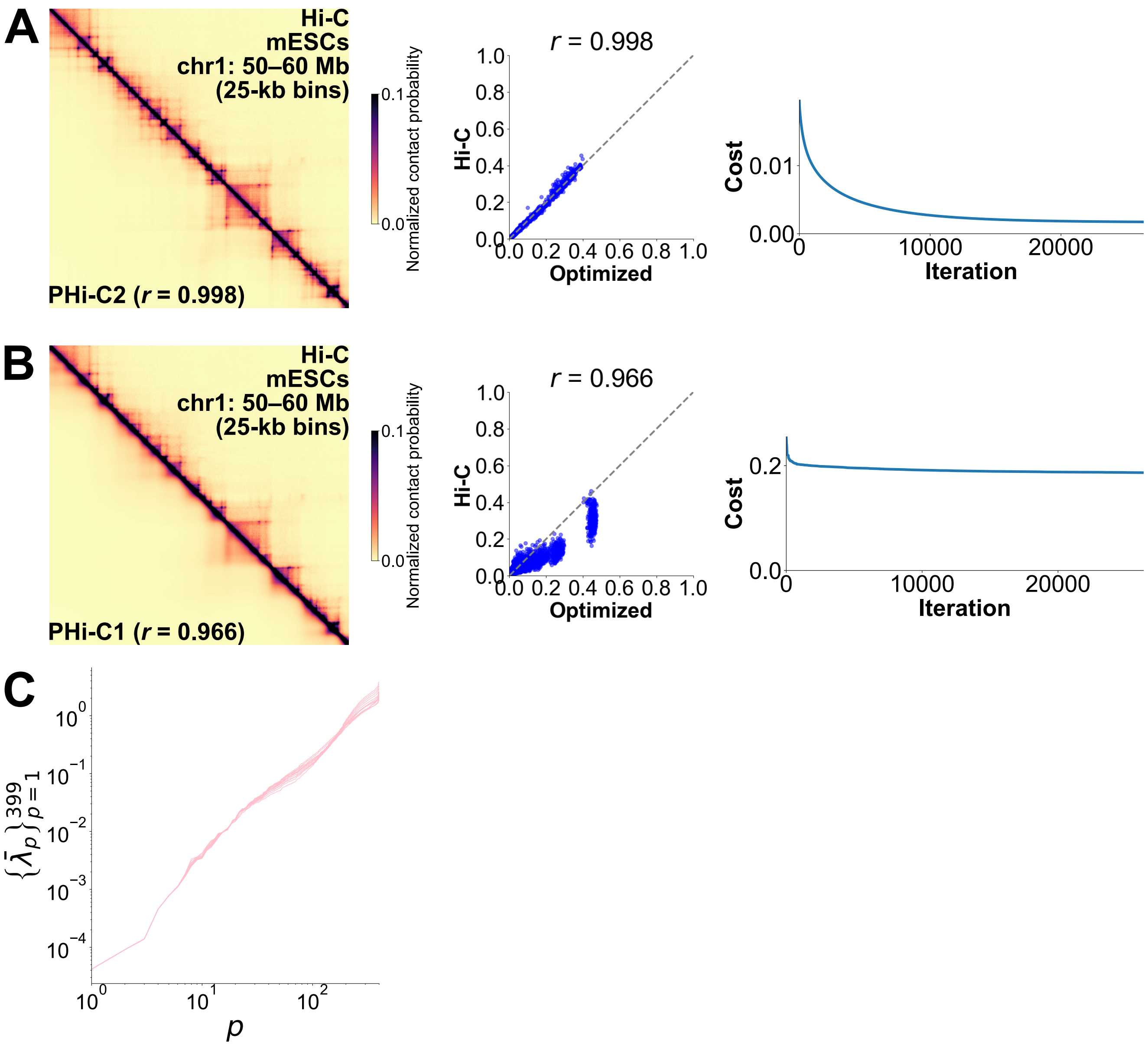


**Supplementary Figure S3. Comparison between (A) PHi-C2 and (B) PHi-C1.** We used Hi-C data of mESCs [chr1: 50–60 Mb (25-kb bins)] as an input. (Left) Heatmap of the contact matrix consists of the input (upper-right) and the optimal (lower-left) matrices. (Middle) Scatter plot with Pearson's correlation coefficient $r$ visualizes the closeness of the input and optimized contact matrices. (Right) Curve of the cost function in the optimization steps. **(C)** Eigenvalues of the induced Laplacian matrix from the optimal solution are positive except for $\bar{\lambda}_0 = 0$. We plotted eigenvalue spectra for the optimal solutions in Supplementary Table S2.

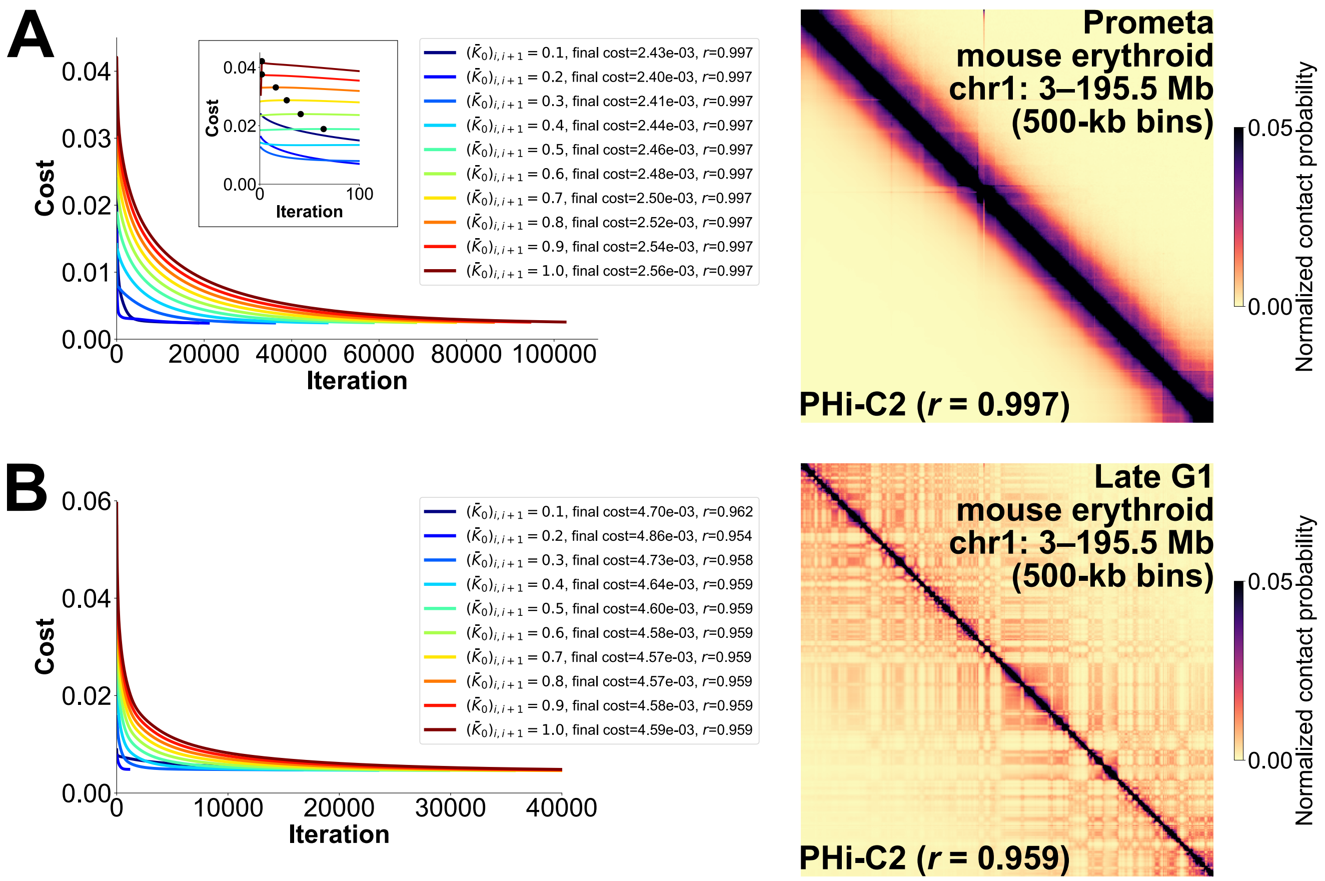


**Supplementary Figure S4. PHi-C2 results for (A) the prometaphase and (B) the late G1 phase.** We used Hi-C data of mouse erythroid cells [chr1: 3–195.5 Mb (500-kb bins)] as an input. (Left) Curves of the cost function. (Right) Heatmap of the contact matrix consists of the input (upper-right) and the optimal (lower-left) matrices. **(A)** (Inset) Magnification in the initial 100 iteration steps. The black points represent the local maximum point for $(\bar{K}_0)_{i,i+1} = 0.5 \sim 1.0$. The heatmap consists of the input contact matrix (upper-right) and the optimal matrix (lower-left) for $(\bar{K}_0)_{i,i+1} = 0.2$. **(B)** The heatmap consists of the input contact matrix (upper-right) and the optimal matrix (lower-left) for $(\bar{K}_0)_{i,i+1} = 0.7$.

#### Supplementary Note S4: Rheology analysis as an additional function

We added the rheology analysis commands as a new function in PHi-C2. We have developed a microrheology method to interpret the Hi-C data as the dynamic 3D genome information[6]. Using the algorithm, we can convert the optimal solution $\bar{K}_{\text{opt}}$ into the rheology spectra of the normalized complex compliance $\bar{J}^*$, the normalized complex modulus $\bar{G}^*$, and the loss tangent $\tan\delta$. Here we show examples for the Hi-C data of mESCs[3] [chr17: 3–95 Mb (250-kb bins), chr17: 41–59.4 Mb (50-kb bins), chr17: 49–52.68 Mb (10-kb bins)] (Supplementary Figure S5). We fixed the matrix size to $368 \times 368$.

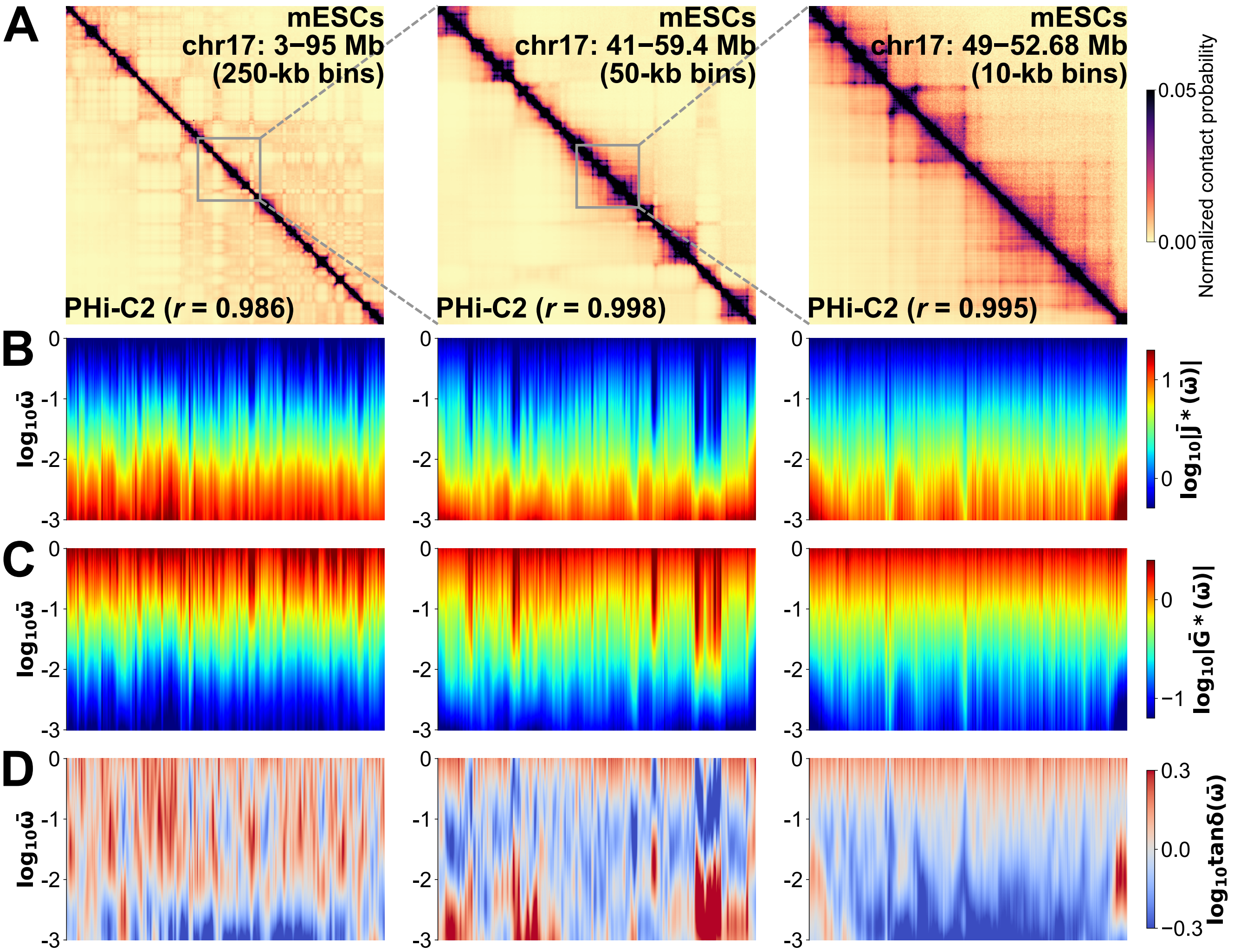


**Supplementary Figure S5. Rehology analysis.** We used Hi-C data of mESCs [(Left) chr17: 3–95 Mb (250-kb bins), (Middle) chr17: 41–59.4 Mb (50-kb bins), (Right) chr17: 49–52.68 Mb (10-kb bins)]. **(A)** Heatmaps of the contact matrix consisting of the input (upper-right) and the optimal (lower-left) matrices. **(B)** Rheology spectra of the normalized complex compliance $|\bar{J}^*(\bar{\omega})|$ as a measure of the dynamic flexibility of three cases. **(C)** Rheology spectra of the normalized complex modulus $|\bar{G}^*(\bar{\omega})|$ as a measure of the dynamic rigidity of three cases. **(D)** Rheology spectra of the loss tangent $\tan\delta(\bar{\omega})$ as a measure of liquid-like ($> 1$) and solid-like ($< 1$) states of three cases.
